# Emergence of a novel interaction between brown bear and cicada due to anthropogenic habitat modification

**DOI:** 10.1101/2020.02.22.960583

**Authors:** Kanji Tomita, Tsutom Hiura

## Abstract

Novel species interactions have generally emerged in ecosystems that are highly modified by human activities. Anthropogenic habitat modification, such as afforestation, is one possible driver of novel species interactions; however, empirical evidence remains scarce. In this study, we show that a novel predator-prey interaction between the brown bear (*Ursus arctos*) and nymphs of a cicada species (*Lyristes bihamatus*) is generated by anthropogenic habitat modification. We evaluated the frequency of brown bear predation on cicada nymphs and the density of cicada nymphs between natural forests and plantations, which are a typical type of human-modified habitat. We found that brown bear predation on cicada nymphs occurred only in the plantations. The density of cicada nymphs in the plantations was significantly higher than in the natural forest. Our results indicate that the plantation leads to the emergence of the bear-cicada interaction due to increasing the density of cicada nymphs. The study draws attention to the overlooked effects of anthropogenic habitat modification on species interactions.

## Introduction

Species interactions vary depending on environmental change. During the Anthropocene, many species now inhabit novel ecosystems characterized by a changing climate, non-native species, and a human-modified habitat [1,2]. In these novel ecosystems, species interactions occur among species that previously never interacted (hereafter: novel interaction) [3–5]. Native species can be susceptible to negative consequences from novel interactions as they lack a co-evolutionary history with their interactors [6,7]. It is difficult to predict when and where novel interactions will emerge, and whether their outcome is positive or negative for native species, as little empirical evidence of novel interactions exist [8,9]. Therefore, understanding the causes and consequences of novel interactions is important for ecosystem management and conservation [8,10].

Most studies on novel interactions focus on interactions between exotic and native species caused by biological invasion [4,11,12]. On the other hand, human-induced environmental changes, such as anthropogenic habitat modification (e.g. land use change) and climate change (e.g. global warming), have been overlooked as a cause of novel interactions, even though these changes alter the behaviour, abundance, and phenology of species, which significantly influences species interactions [13–15]. Moreover, these environmental changes provide consumers with opportunities for acquiring novel resources [8,16]. However, to our knowledge, there are no studies showing empirical evidence that anthropogenic habitat modification generates novel interactions among native species (but suggested by Fagan et al. [17]).

We focus on a predator-prey interaction between the brown bear (*Ursus arctos*) and nymphs of a cicada species (*Lyristes bihamatus*) in the Shiretoko World Natural Heritage (hereafter: SWH), northern Japan. This is a case of a novel interaction between native species because both species are native but brown bears have only started preying on cicada nymphs since 2000 in the area [18]. The interaction is generated by a driver that is not associated with the invasion of either predator or prey species. The plantation, a typical type of human-modified habitat [19], is one possible driver of the bear-cicada interaction, given brown bear predation on cicada nymphs were frequently observed in larch (*Larix kaempferi*) plantations [18].

In this study, we evaluated the frequency of predation and the density of cicada nymphs between the natural forest and the plantations. We made the following predictions according to our previous findings [18]: (1) brown bear predation on cicada nymphs occurs more frequently in the plantations than in the natural forest; (2) the density of cicada nymphs is higher in the plantations than in the natural forest; (3) there is a positive relationship between predation frequency and the density of cicada nymphs.

## Materials & Methods

### (a) Site Description

The study was conducted in forests located on the western parts of the Shiretoko Peninsula (Fig. S1). The elevation ranged from 120 m to 220 m. UNESCO has certified the area as a World Heritage site because it represents one of the richest northern temperate ecosystems in the world (http://whc.unesco.org/en/list/1193). The natural forests are conifer-broadleaved mixed forests, mainly consisting of Sakhalin fir (*Abies sachalinensis*) and Mongolian oak (*Quercus crispula*). Natural forests account for 82% of the forest area in the study site, while plantations account for 18% of the total forest area. Spruce (*Picea glehnii*), larch and fir plantations account for 13%, 4%, and 1% of the total forest area, respectively. Most of the larch and fir plantations were established during the late 1970s, whereas the spruce plantations were established during the early 1990s [20]. The vegetation map of the study site is shown in Fig. S2.

The SWH has one of the highest densities of brown bear in the world[21]. Within the study site, food items of the brown bears change across seasons; that is, herbaceous plants are consumed in spring, herbaceous plants, ants and cicada nymphs are consumed in summer, and *Q. crispula* acorns and anadromous salmon are consumed in autumn [18,22,23]. Within the study area at least 11 individual bears were preying on cicada nymphs, including two subadults, two solitary female adults, and three females with cubs [18]. Two native cicada species, *Lyristes bihamatus* and *Terpnosia nigricosta*, occur in the SNH and emerge during late summer and spring to early summer, respectively. In the study site, brown bears prey on the nymphs of *L. bihamatus*, but not *T. nigricosta* [18]. Hence, this study focuses on *L. bihamatus* as a prey item of bears and the term “cicada” refers to *L. bihamatus*.

### (b) Field Survey

From late August to September 2018, 100 m^2^ survey plots were set in the following forest types: larch plantations (N = 15), fir plantations (N = 12), spruce plantations (N = 15), and natural forests (N = 30). The location of the survey plots is shown in Fig. S2. A larger number of plots were in the natural forest, as this is the highest proportion of forest for all forest types. Stand characteristics of each forest type are shown in Table 1. The density of cicada nymphs was measured using the density of cicada exuviae collected from all trees (diameter breast height, DBH > 2 cm) within the plots. Sampling heights of trees were less than 3 m. Brown bear predation on cicada nymphs was measured by observing digging traces of brown bears, as the bears dig up soil when preying on cicada nymphs [18]. According to our preliminary observations by camera traps, brown bears usually dig for cicada nymphs near the base of a tree. We evaluated the predation frequency per each plot as the proportion of trees that had digging traces within a 50 cm radius from the base of a tree for all trees (DBH > 2 cm) in the plot.

**Table 1.** Stand characteristics across the forest types.

| Forest type(Number of plots) | Dominant<br>tree species (%Basal area) | Density of trees<br>(Trees 100m<sup>-2</sup>) | Basal area<br>(m<sup>2</sup> 100m<sup>-2</sup>) |
| --- | --- | --- | --- |
| Larch plantation (N=15) | *Larix kaempferi* (97%) | 32.06 ± 7.78 | 0.12 ± 0.04 |
| Fir plantation (N=12) | *Abies sachalinensis* (84%) | 30.17 ± 10.80 | 0.11 ± 0.04 |
| Spruce plantation (N=15) | *Picea glehnii* (95%) | 27.14 ± 6.36 | 0.09 ± 0.03 |
| Natural forest (N=30) | *Abies sachalinensis* (31%) | 30.87 ± 10.73 | 0.10 ± 0.05 |
|  | *Quercus crispula* (28%) |  |  |

### (c) Statistical Analysis

Generalized linear models (GLMs) with a log link, Poisson error distribution and Tukey post hoc tests were used to examine the differences in predation frequency and the density of cicada nymph among the forest types. When the GLM indicated a significant difference (*p-value* < 0.05) of one forest type from others, we performed a multiple comparison between the forest types. In the GLM analysis for predation frequency, we introduced an offset term as the number of trees (log-transformed) to adjust for differences in the number of trees among the survey plots. To examine the effects of cicada nymph density on predation frequency, we used GLMs with log links and Poisson error distributions for each forest type. Number of trees (log-transformed) were included as an offset term in the GLMs. All statistical analyses were conducted in R version 3.5.1 [24].

## Results

The GLMs indicated a significant effect of forest type on predation frequency and the density of cicada nymphs. Surprisingly, brown bear predation on cicada nymphs only occurred in plantation plots, not the natural forest plots even in which mainly composed of fir species (Fig. 1A). The density of cicada nymphs was lowest in the natural forest plots (Fig. 1B). The predation frequency was highest in the larch plantation plots, but the density of cicada nymphs did not differ from in the fir plantation plots, which had a lower predation frequency (Fig. 1). The spruce plantation plots had a lower predation frequency and the density of cicada nymphs than other types of plantation plots (Fig. 1). The density of cicada nymphs positively affected the predation frequency in all plantation types (GLM, p < 0.05 Fig. 2), suggesting that predation occurred more frequently as the density of cicada nymphs increased.

**Figure 1.**
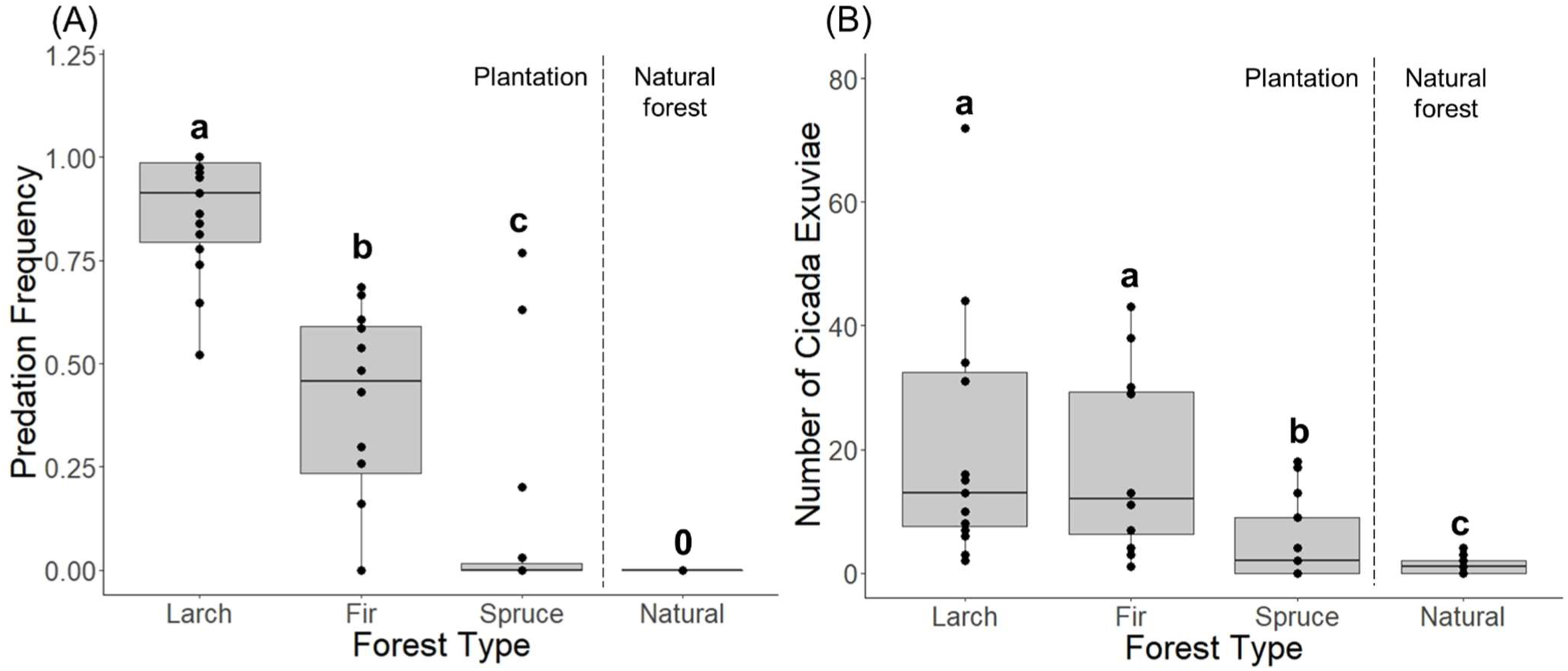
(A) Frequency of predation of brown bear on cicada nymphs and (B) the density of cicada nymphs in larch plantations, fir plantations, spruce plantations, and natural forests. Different letters indicate significant differences (Tukey test, p < 0.05).

**Figure 2.**
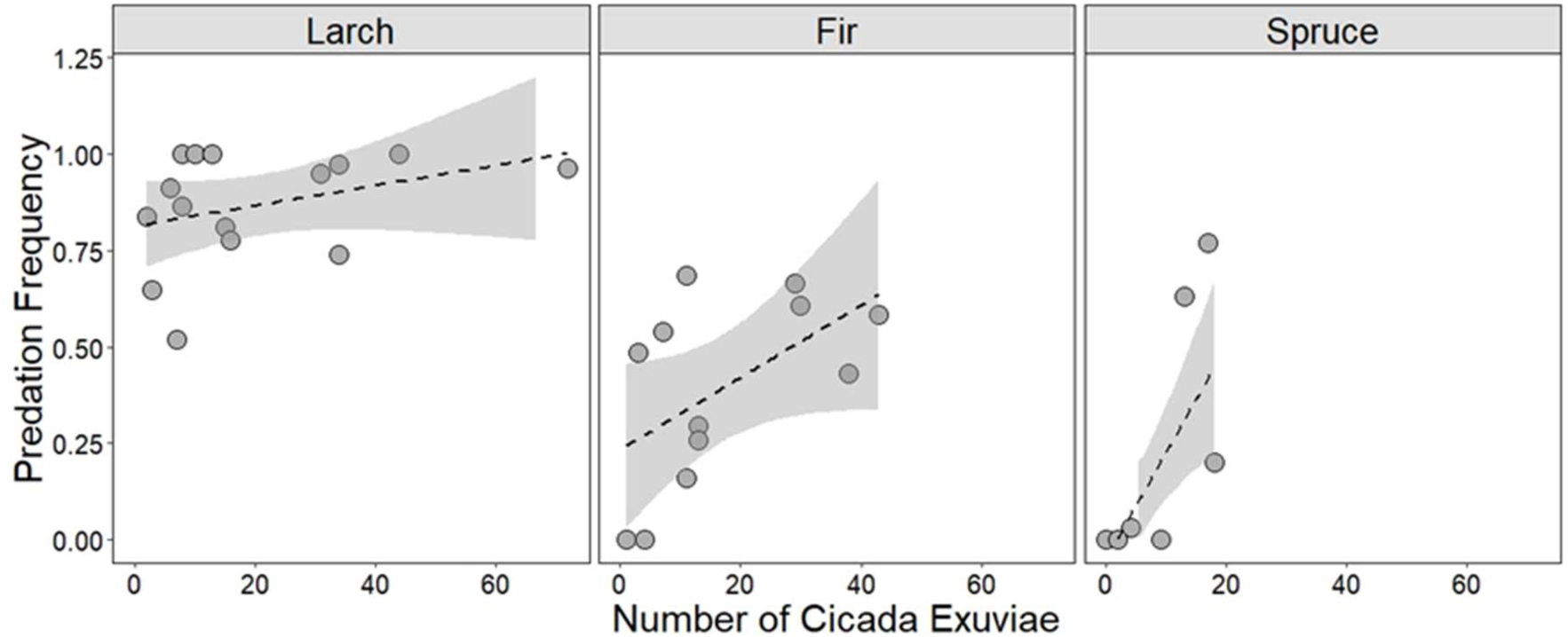
Relationships between the predation of brown bear on cicada nymphs and the density of cicada nymphs. Dashed lines show the linear model predictions with shaded areas indicating the 95% CI. The relationships are significantly positive in larch plantation (Left), fir plantation (Center), spruce plantation (Right) (GLM, p < 0.05).

## Discussion

The results of our study generally supported the predictions. In particular, brown bear predation on cicada nymphs only occurred in the plantations, not the natural forests. Furthermore, in the SWH, the plantations were established during the late 1970s, with trees reaching maturation around 2000 [20,25]. Because cicada nymphs generally grow under mature trees [26], they could inhabit the plantations since 2000 in this area. These indicate the high density of cicada nymphs have occurred in the plantation since about 2000; consequently, the bear-cicada interaction has emerged. To our knowledge, this is the first study presenting empirical evidence of a novel interaction between native species due to anthropogenic habitat modification.

Our results indicate that the differences in predation frequency among forest types can generally be explained by the density of cicada nymphs (Figs. 1,2). However, the predation frequency in larch plantation was higher than in fir plantation despite no differences in the density of cicada nymphs between these types of plantations. The most plausible reason for this is that the density of cicada nymphs in the larch plantations was underestimated. The use of cicada exuviae as a proxy for the nymph density may have resulted in an underestimation of actual nymph density because it does not account for nymphs that were already preyed upon by the brown bears. Another reason is that brown bears had fewer chances of encountering fir plantations than larch plantations because the total area of larch plantations was larger than fir plantations at the study site (Fig. S2). One final reason is the social transmission of information from mothers to their cubs that cicada nymphs were abundant in larch plantations, which might constrain brown bears to prey on nymphs within the larch plantations. Tomita & Hiura [18] showed that bears preying on cicada nymphs mainly consisted of solitary adult females and females with cubs. Since socially learned foraging behaviours in bears are expected to be female biased [27,28], the predatory behaviour might propagate through the brown bear population via social learning. Moreover, because brown bears in the SWH show female-biased philopatry [29], female bears learning the behaviour may stay within the study site.

Consistent with our results, other studies have shown high densities of cicada nymphs in human-modified habitats, such as forest edges and plantations [30–32], though the reasons are unclear (but see Yang [33]). This suggests that anthropogenic habitat modifications provide high quality habitat for cicadas. Since some birds and mammals prey on cicadas [18,34,35], anthropogenic habitat modifications can positively affect these predators via increasing availability of cicada nymphs.

Although the density of cicada nymphs in the larch, fir, and spruce plantations was underestimated compared to natural forests due to predation by brown bears, the density in these plantations was still significantly higher than in the natural forests (Fig. 1B). This indicates the positive effects of plantations on the cicada population compensate for the negative effects of increased predation. Therefore, the bear-cicada interaction does not seem to lead to negative consequences for the cicada population, a notion supported by the fact that the interaction has continued for 20 years at the study site [18].

Detecting novel interactions among native species is challenging because it can be difficult to determine whether they are novel or are pre-existing interactions that have been overlooked. Therefore, we might be failing to notice how anthropogenic habitat modifications generate novel interactions among native species, even though the impacts of anthropogenic habitat modification on native species assemblages continues to strengthen around the world [36]. Our study draws attention to an overlooked aspect of the effects of anthropogenic habitat modification on species interactions. Furthermore, our results showed that this bear-cicada interaction does not seem to have negative consequences for the cicada population. On the other hand, Liu et al. [37] showed how global warming caused a novel interaction between a native herbivore and a native plant that had significant negative impacts on plant reproduction. Further studies are needed to understand the consequences of novel interactions caused by anthropogenic environmental changes, such as habitat modification and climate change.

## Supporting information

Supplemental Meterial Fig.S1 and Fig.S2

## Acknowledgments

We thank members of Shiretoko Nature Foundation for providing information on study site. We also thank H. Maita and Daisetsu I. and T. Itoh for field survey assistance and I. Koizumi for providing valuable comments on the manuscript, and I. Tanada for advice the English of the manuscript.

