## Supplemental Meterial Fig.S1 and Fig.S2 for "Emergence of a novel interaction between brown bear and cicada due to anthropogenic habitat modification"

### Supplemental Material

Figure S1 (A) Map of Shiretoko peninsula, northern Japan. (B)Map of Shiretoko peninsula, including the study site.

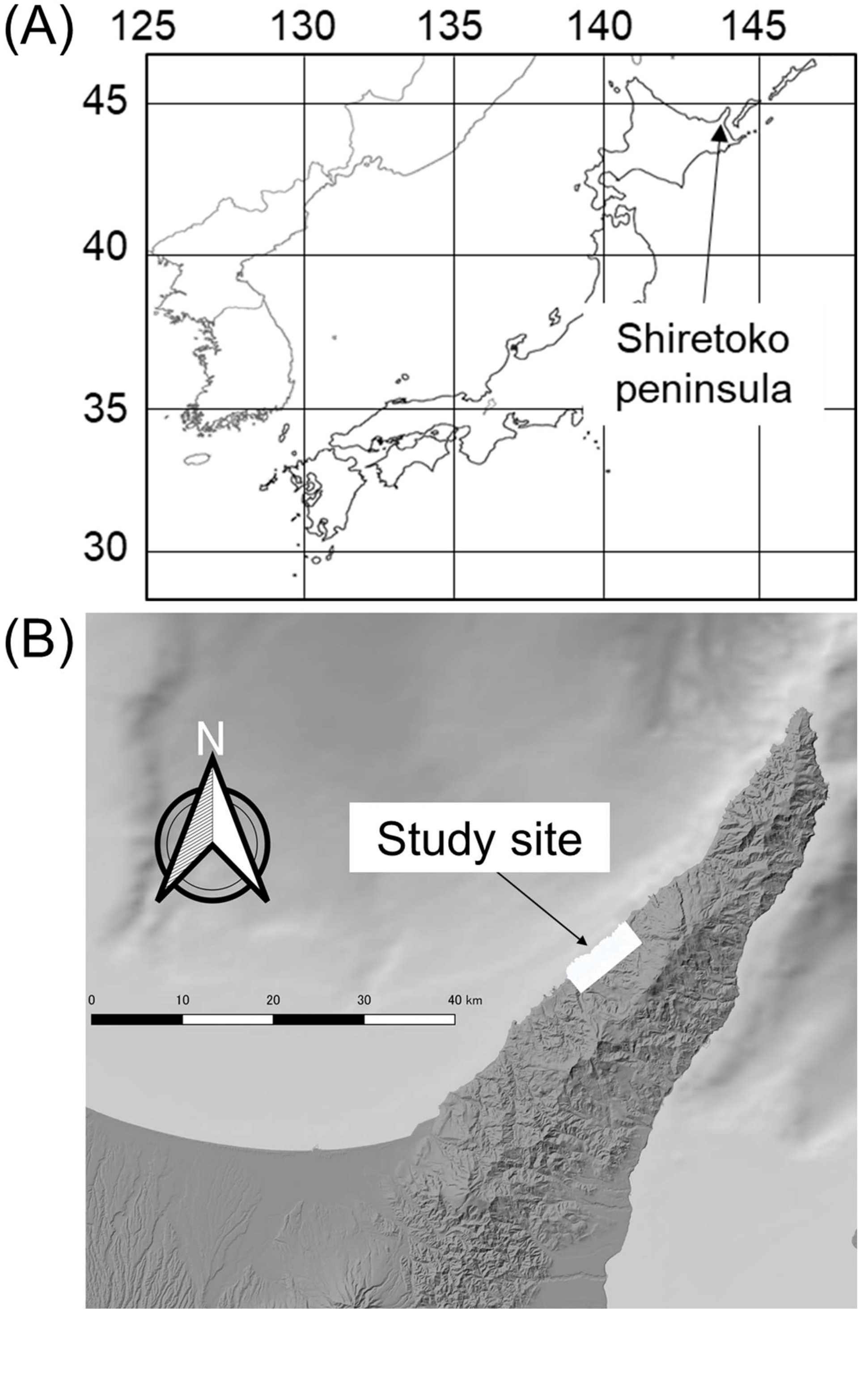

Figure S2 Location of the survey plots in vegetation map of the study site.

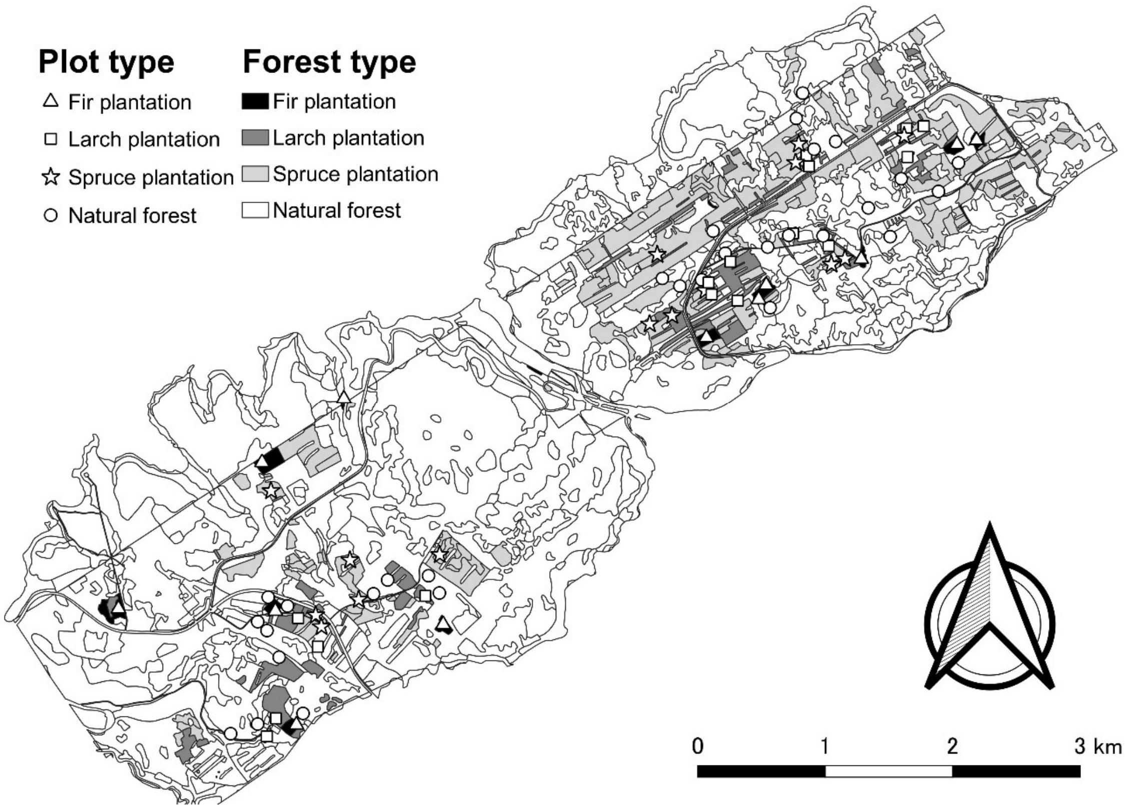
